# The effect of estimation time window length on overlap correction in EEG data

**DOI:** 10.1101/2023.06.05.543689

**Authors:** René Skukies, Benedikt V. Ehinger

## Abstract

Regression Event-Related-Potentials (ERPs) with overlap correction (also referred to, as linear deconvolution, or temporal response functions) are becoming more popular for the analysis of Electroencephalography (EEG) data. A common question for the analyst is, how to specify the length of the estimation windows. Long estimation windows might capture all relevant event-related activity, but might introduce artifacts due to overfit, short estimation windows might not overfit, but also might not capture all (overlapping) activity, and thereby introduce bias.

Using a systematic simulation approach, we show that longer rather than shorter time windows should be preferred for typical EEG designs. We further provide an interactive app to visualize various design parameters: https://estimationwindow.ccn2023.s-ccs.de

## Introduction

Neural activity is inherently noisy. To attenuate this noise, data is often time locked to an event and averaged over many repetitions. Assuming that the noise is uncorrelated to the event onsets and given enough repetitions, this approach will dampen the noise and recover the underlying signal.

However, when using averaging, we, often implicitly, assume that brain activity of adjacent events does not overlap in time. To give an example where this assumption is violated, let’s imagine a decision-making experiment where participants have to decide if a stimulus belongs to one of two categories. Typically, such findings are interpreted in light of evidence accumulation models (Pisauro, Fouragnan, Retzler, & Philiastides, 2017), where activity for fast trials shows a steeper increase in activity time locked to the button press, compared to slower trials. An alternative explanation is based on temporal overlap of ERPs: In fast trials, stimulus onsets and button presses are closer together in time compared to slow trials, and both stimuli and button presses elicit brain activity, often persisting longer in time than the actual distance of events. Indeed, we recently showed that the CPP-related evidence accumulation activity observed by others, can be explained by an overlap-corrected model (Frömer, Nassar, Ehinger, & Shenhav, 2022). This highlights one paradigm, where appropriate overlap correction is necessary and simple averaging is not enough.

The most common approach to model such overlap is based on linear deconvolution models (Smith & Kutas, 2015), commonly applied to event-related fMRI experiments (Dale & Buckner, 1997). In a nutshell, instead of modelling time-locked data, the continuous EEG is modelled as a mixture of overlapping ERPs. It is assumed that each EEG sample can be modelled by the summation of the ERPs of the overlapping events. In practice, we define a linear model incorporating all relevant information about each of our events, resulting in one design-matrix per event type. Next, we define **estimation windows (EW)** around all event onsets, and “time-expand” our design-matrices around each event onsets, resulting in a much taller and wider design-matrix — taller to cover all EEG samples; and wider to reflect that we estimate the whole ERP of all events concurrently (in contrast to the time-point by time-point operations when using averaging or mass-univariate analyses)

In the classical averaging analysis, we also make use of EWs - but critically, the actual EW length does not influence the estimate at other time-points. In striking contrast, choosing the correct EWs for deconvolution analysis is crucial. Indeed, the question of choosing EWs is one of the most common questions we get with collaborators or at EEG workshops.

In this paper, we investigate the influence of EW length. Too short EWs will not be able to capture the overlapping signals correctly, too long EWs could lead to overfit. To do so, we simulate ERPs, overlap them in time, and try to recover the original shape while varying the lengths of the EWs.

## Methods

Using the UnfoldSim.jl toolbox (Ehinger & Lips, 2023), we generated proto-ERPs of 0.450 s length, based on scaled Hanning windows at 250Hz, resembling a typical P1-N1-P3 complex (Figure 1, gray line). We simulated 400 event-onsets, where the event-distances were sampled from a uniform distribution from 0.25-0.35s. The continuous EEG was assembled, and we added pink noise to the signal (single trial signal-to-noise at component peak ≈ 0.6).

**Figure 1:**
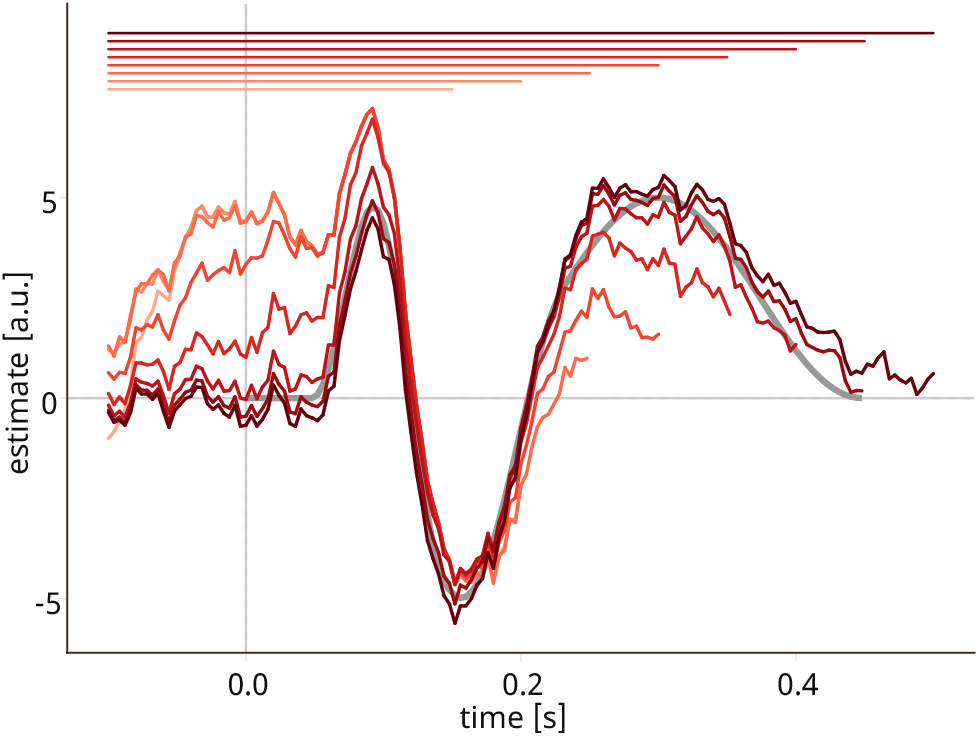
Overlap corrected ERPs against different EW durations, one simulation. Gray line shows ground truth. Red lines show EWs for durations of [0.15:0.05:0.50] (c.f. indicator bars)

Using linear deconvolution, we then tried to recover the proto-ERP using the Unfold.jl toolbox (Ehinger & Dimigen, 2019; Ehinger et al., 2023). To investigate the influence of EW size, we analyzed the same simulated data using eight different window sizes (all EWS started at -0.1s, ranging to 0.1:0.05:0.5s or 3s). The overlap-corrected responses were then compared to the ground truth by calculating the mean squared error (MSE) between 0 and 0.45s. This made it necessary that EWs shorter than the ground truth were zero-padded to match the length of the ground truth, and EWs longer than the ground truth were truncated. Such an approach potentially biases against short time-windows (as overfit outside the ground truth time-window is not considered). Thus, we repeated the analysis with the respective “full”-MSE for each EW.

This procedure was repeated a total of 100 times with different random seeds.

## Results

To show the influence of estimation window on linear deconvolution analysis, we systematically vary the EW size on simulated data.

As visible in Figure 1 for a single simulation, EWs that are shorter than the ground-truth activity show a systematic bias away from the ground truth. Indeed, in a quantitative analysis (Figure 2) we see a near linear decline of MSE with increasing EW until the EWs match the ground truth time window (short EW 0.15s, mean MSE: 10.44, SD: 1.19 — ground-truth matching EW: 0.45s, mean MSE: 0.46, SD: 0.38).

**Figure 2:**
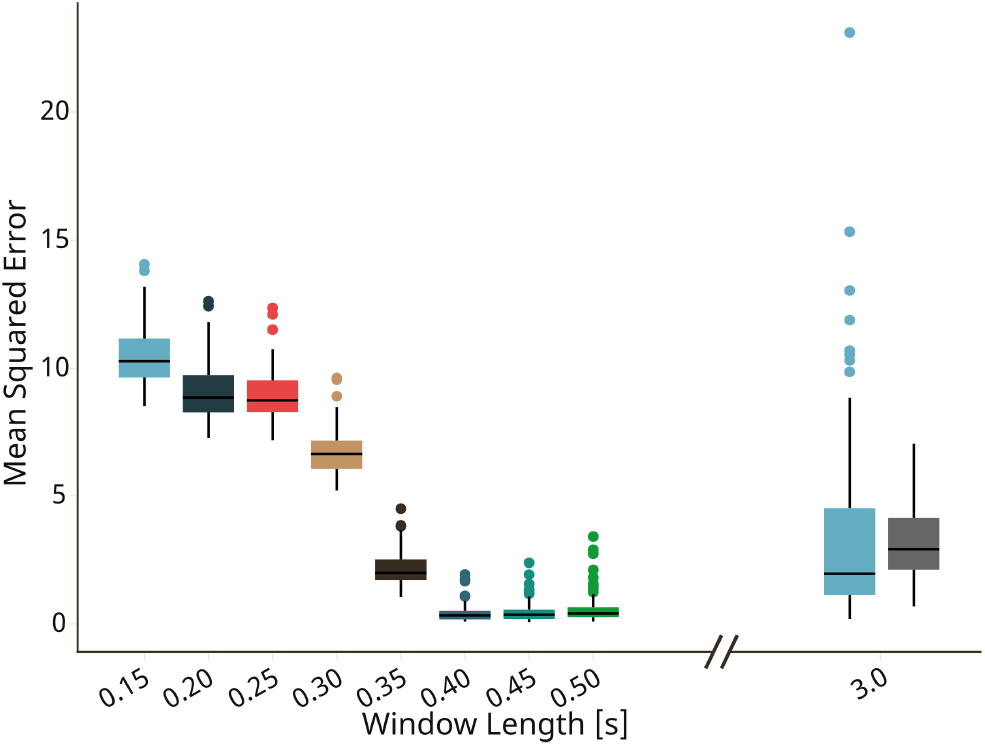
MSE across varying EWs during the ground truth time window, except for the last gray boxplot, which shows the 3s-EW MSE over the whole 3s

Increasing the EW beyond the ground truth time window resulted in worse MSE (EW: 0.5, mean MSE: 0.59, SD: 0.58), which emphasized in much longer EWs (EW: 3.0s, mean MSE: 3.69, SD: 3.69), even though in this analysis, we evaluate MSE in the period of the ground truth time window alone.

As indicated above, only testing the MSE in the ground truth time window underestimates the overfit of the total estimated ERP. We thus repeated the analysis of the long time-windows and calculated the MSE in their respective EWs, effectively zero-padding the ground truth. As seen in Figure 2 (gray box-plot) MSE values along the entire EW length only worsened slightly, and we observe a decrease in variance. This likely reflects the decreasing influence of the initial ground truth activity against the longer lasting baseline period. In other words, for long enough EW duration, MSE approaches the pure noise variance. However, investigating the results from just a single iteration (Figure 3) we can see that estimates using long EWs are strongly contaminated by noise, potentially misleading the researcher.

**Figure 3:**
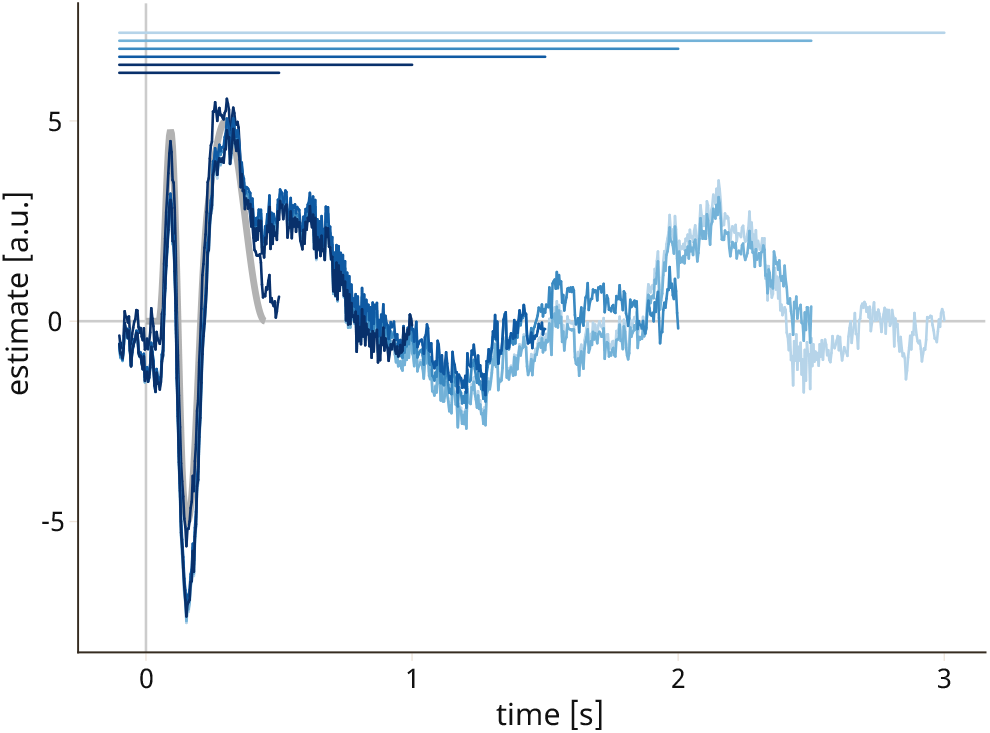
Same as Figure 1, but with longer EWs. Grey line shows ground truth. Blue lines show EWs for durations of [0.5:0.5:3.0] (c.f. indicator bars)

Finally, we round off our analysis with several robustness analyses: The main pattern depicted in Figure 2 persisted regardless of noise (**PinkNoise**, RedNoise, WhiteNoise), signal-to-noise (1, **0.6**, 0.25), or proto-ERP shape (**P1-N1-P3**, P1-N1-N3, P1-N1, P3-only).

## Discussion

Choosing the estimation time window length is critical for analyses relying on overlap correction. Our simulations show that the best performing models are those with an EW length which matches (or closely matches) the actual underlying activity.

For analyzing real-world EEG data it follows that researchers should — a priori — make an educated guess about the length of the underlying EEG activity and select this as their EW. This also suggests to use event windows with different sizes between events (as possible in e.g. Unfold.jl (Ehinger et al., 2023)). Given that the overfit is only of moderate size, when choosing longer time-windows, we further recommend to generally err on the longer side.

Lastly, two more things to keep in mind :

1. modifying the EW after already previewing a model fit might not be as inconsequential as in the classical analyses. In the strictest sense, such a post-hoc parameter change can lead to overfit, as the analysis parameter was informed by the data itself. We are not aware of studies showing the extent of harm that such an overfit/circular procedure has, if any. In case one pre-specified a time-window not capturing the activity, we recommend prolonging the EW and to report this change transparently in your manuscript.
2. In practice, we have observed in several datasets a lingering activation beyond 1s, sometimes approaching 0μV only after the highpass-filter enforced it. Non-systematic tests showed little to no influence on the more immediate ERP-shape when truncating this low-μV activity.

Summarized, using a simulation study, we address the gap on how to specify the estimation time window when using linear deconvolution analyses in EEG data.

## Acknowledgments

Funded by Deutsche Forschungsgemeinschaft (DFG, German Research Foundation) under Germany’s Excellence Strategy – EXC 2075 – 390740016. We acknowledge the support by the Stuttgart Center for Simulation Science (SimTech). All the necessary code for the analysis can be found under http://doi.org/10.5281/zenodo.7785501.

## Package References

**Julia 1.8.3** (Bezanson, Karpinski, Shah, & Edelman, 2012) **- AlgebraOfGraphics.jl 0.6.14** - **CairoMakie.jl 0.10.2** (Danisch & Krumbiegel, 2021) - **DataFrames.jl 1.5** - **StatsBase.jl 0.33.21** (Lin et al., 2022) **- Pluto.jl 0.19.22** (van der Plas et al., 2023) - **UnfoldSim 0.1.2** (Ehinger & Lips, 2023) **- Unfold.jl 0.3.13** (Ehinger et al., 2023)

